# Otx2 stimulates adult retinal ganglion cell regeneration

**DOI:** 10.1101/2020.10.06.327999

**Authors:** Raoul Torero-Ibad, Nicole Quenech’du, Alain Prochiantz, Kenneth L. Moya

## Abstract

Retinal ganglion cell axons provide the only link between the light sensitive and photon transducing neural retina and visual centers of the brain. Retinal ganglion cell axon degeneration occurs in a number of blinding diseases and the ability to stimulate axon regeneration from surviving ganglion cells could provide the anatomic substrate for restoration of vision. OTX2 is a homeoprotein transcription factor expressed in the retina and previous studies showed that, in response to stress, exogenous OTX2 increases the *in vitro* and *in vivo* survival of retinal ganglion cells. The present results show that, in addition to promoting adult retinal ganglion cell survival, OTX2 also stimulates the regeneration of their axons *in vitro* and *in vivo*. This dual activity of OTX2 on retinal ganglion cell survival and regeneration is of potential interest for degenerative diseases affecting this cell type.

## Introduction

Neurodegenerative diseases that affect retinal ganglion cells (RGCs) lead to a degradation of vision and, very often, to blindness. In diabetic retinopathy, optic neuropathies, and glaucoma, RGC axons degenerate and the cell bodies die. Considering just glaucoma, there are an estimated 1 million people afflicted in France (source https://www.aveuglesdefrance.org/glaucome accessed 28-07-2020) with direct medical costs of 455-969€/patient/year in Europe depending on severity (Traverso et al., 2005). Thus, any factor that might protect against RGC loss and stimulate the regrowth of damaged RGC axons would be of significant therapeutic interest in multiple diseases of vision.

Homeoprotein transcription factors are also morphogens with the capacity to transfer between cells and to regulate gene transcription, protein translation as well as chromatin remodeling in a non-cell autonomous way (see (Di Nardo et al., 2018) for review). Among this family of transcription factors, OTX2 is essential at many steps of eye development, including the differentiation of photoreceptors and bipolar cells (Koike et al., 2007; Martinez-Morales et al., 2001; Muranishi et al., 2011; Nishida et al., 2003). In the adult, OTX2 remains expressed in the two latter cell types and its reduced activity in hypomorph mice results in progressive loss of photoreceptors and bipolar cells starting at around 2 months of age (Bernard et al., 2014). In RGCs the *Otx2* locus is silent, but the protein can be detected in ganglion cell layer cells in both mouse and human retina suggesting that RGCs take up the protein, most likely from bipolar cells (Azzolini et al., 2013; Sugiyama et al., 2008).

Previously, it was reported that exogenous recombinant OTX2 promotes the survival of damaged adult RGCs *in vitro* and *in vivo* (Torero Ibad et al., 2011). In cultures of dissociated adult mouse retinal cells, OTX2 increased RGC survival up to over four-fold compared to untreated cultures. Cultures of purified adult rat RGCs showed that OTX2 acts directly on RGCs to promote their survival. Intraocular injection of NMDA rapidly kills RGCs and is considered as an acute model of glaucoma. When co-injected with NMDA, OTX2 provided full protection against excitotoxic RGC loss. In addition, OTX2 protected against the loss of visual acuity caused by NMDA injection.

In preliminary *in vitro* experiments, at higher concentrations of OTX2, the number of RGCs with an axon and the length of regenerated axons seemed to be increased. Here we directly examine and quantify OTX2 effects on adult RGC axon regeneration *in vitro* and *in vivo*. The results show that exogenous OTX2 stimulates the regrowth of axons from RGCs in cultures of dissociated adult retinal cells and from explants of adult retinal tissue and that RGCs respond directly to OTX2 as regrowth is observed in cultures of purified adult rat RGCs. Importantly, after nerve crush, we observe a positive effect of OTX2 on the number of regenerating axons up to the optic chiasm within 14 days post crush and a very modest level of acuity absent in control mice.

Combined with published results (Bernard et al., 2014; Torero Ibad et al., 2011), the present study further demonstrates the interest of OTX2 for promoting the survival of injured adult RGCs and reveals its potential for stimulating RGC axon regeneration, suggesting that it may be used in ocular diseases in which RGCs are lost such as glaucoma and optic neuropathies.

## Methods and Materials

### Preparation of recombinant OTX2

Recombinant mouse OTX2 was produced and purified as described (Torero Ibad et al., 2011). Immunofluorescence for neurofilament NF200 (1:500, Sigma, catalog #N4142) and anti-rabbit Alexa Fluor 545 (Invitrogen).

### Culture of purified adult RGCs

Immunopurified RGCs were prepared from adult rat retinas based on (Fuchs et al., 2005). Eight-week-old Long-Evans rats were killed by CO2 and cervical dislocation, the eyes removed and retinas dissected in CO2-independent media as above. Retinal tissue was dissociated with papain and mechanical disruption. RGCs were captured by substrate bound anti-Thy1 (hybridoma supernatant TIB-103, LGC Promochem). And then plated as for the mixed cultures. OTX2 or vehicle was added at the time of plating and cells maintained for six days at 37°C in 5% CO2. Live cells were incubated with 2 µM calcein acetoxymethyl ester (Thermo Fischer) for 1 h at 37°C, and the number of living cells and the number of cells with a neurite longer than two soma diameters on six coverslips per condition were counted on an epifluorescence microscope with a 488 nm filter. The investigator counting was blind to the treatment condition.

### Retinal explants

Eight-week-old C57 Bl6 mice or Long Evans rats were killed by CO2 and cervical dislocation. Eyes were removed and retina flattened RGC side up on mixed ester cellulose filters (Whatman, 10409770). One mm^2^ explant squares were cut with a McIlwain tissue chopper and the explant/filter squares were placed RGC side down on poly-D-lysine/laminin coated coverslips and incubated in supplemented Neurobasal A Neurobasal A (Thermo Fischer) complemented with B27 serum-free supplement (Thermo Fischer), 500 mM glutamine, 25 mM glutamate, 25 mM aspartate, and antibiotic/antimycotic. OTX2 or vehicle was added at the time of plating. At 6 DIV, explant cultures were fixed in buffered 4%PFA and labeled with anti-NF 200 as above. Explant images were acquired with Nikon 90i microscope, opened in Fiji v 2.0.0-rc-15/1.49k and axon/axon fascicles emanating from the explant manually traced by an investigator blind to the treatment condition. Total axon/fascicle length per explant was determined with the measure command and averaged for at least 4 explants per condition.

### Optic nerve crush

Optic nerve crush (ONC) was performed as described (de Lima et al., 2012). In brief, mice were anesthetized with xylazine/ketamine (Imalgène 500 Virbac France, 100 mg/kg, Rompun 2% Bayer, 10 mg/kg). Tropicamide (Mydriaticum 0.5% Théa, France) was used to dilate the iris. The right eye was partially exorbited and the conjunctiva was cut with irretectomy scissors along the ora serrata starting at the dorsal pole. The superior rectus was cut proximal to the *limitans*, the optic nerve adjacent to the globe was exposed by blunt dissection and crushed under visual guidance for 10 secs using 5/45 45°-angled jeweler’s forceps (WPI). The crush site was visually verified by two investigators, one using the surgical scope and the other via live imaging on a video screen. The eye was repositioned in the orbit and irrigated with normal saline. Within 5-15 min OTX2 (30-205ng in 1-2µl) or vehicle was injected intraocularly as described (Torero Ibad et al., 2011). After eye injection, the mice were placed in a heated chamber at 37°C until recovered from the anesthesia and returned to their home cages

### Anatomical analyses

Two weeks after OTX2 injection, mice were transcardially perfused with PBS followed by 4% PFA, eyes removed and the dissected retina prepared with four cuts for flat mount processing and post fixation in 4% buffered PFA. Optic nerves were dissected from posterior to the eye to the optic chiasm, post fixed in 4% PFA on a paper filter and prepared for cryosection. Free-floating retinas were processed for Brn3a (Santa Cruz antibodies, sc-31984) immunofluorescence and RGCs counted in eight fields of view per retina using analyze particles function on Fiji software (Danias et al., 2002). Optic nerves were processed using anti-GAP-43, a marker for regrowing axons (Leon et al., 2000).

### Visual acuity testing

Mice were tested for visual acuity a week before ONC, three weeks after ONC and at 14 weeks after ONC. The left, non-affected eye was sutured shut, one day prior acuity testing. For suturing, the mice were anesthetized with xylazine/ketamine as above, the skin around the eye swabbed with 70% ethanol, avoiding to contact the eye and the lid sutured closed with two sutures of 6/0 silk. Visual acuity was tested using the OptomotryTM. Mice were placed on an elevated platform in an arena surrounded by four high definition video screens. Black and white 100% contrast square-wave gratings of different spatial frequencies were presented in a semi-shuffled order. In the optokinetic response, mice make head turns in the temporo-nasal direction of the stimulus direction. If the stimulus is clockwise, the head turn is from left to right and driven by the left eye. If the stimulus is counterclockwise, the head turn is from right to left and driven by the right eye. The experimenter was blind to spatial frequency and to the direction of movement presented. If no head turn was signaled within 2 min, a pattern of lower spatial frequency was presented. If a head turn was observed, the next higher spatial frequency was presented to establish the acuity threshold.

All animal procedures were carried out in accordance with French national and European directives for the care and use of research animals and approved by the local “*Comité d’éthique en expérimentation animale n°59*” and authorization n° 00702.01 delivered by the French “*Ministère de l’enseignement supérieur et de la recherche*”.

## Results

RGCs were purified from adult rat retina and OTX2 stimulated RGC survival in a bell-shaped dose-response curve, independently corroborating previously reported results with a significant 2-fold increase at 3.3nM Otx2 (Figure 1A). Importantly, the number of live RGCs with neurites was also significantly higher by about 3-fold at 3.3 nM exogenous OTX2 compared to controls (Figure 1B). Comparing the number of live RGCs and those with neurites, OTX2 stimulated neurite growth in about one-fourth of the surviving RGCs.

**Figure 1.**
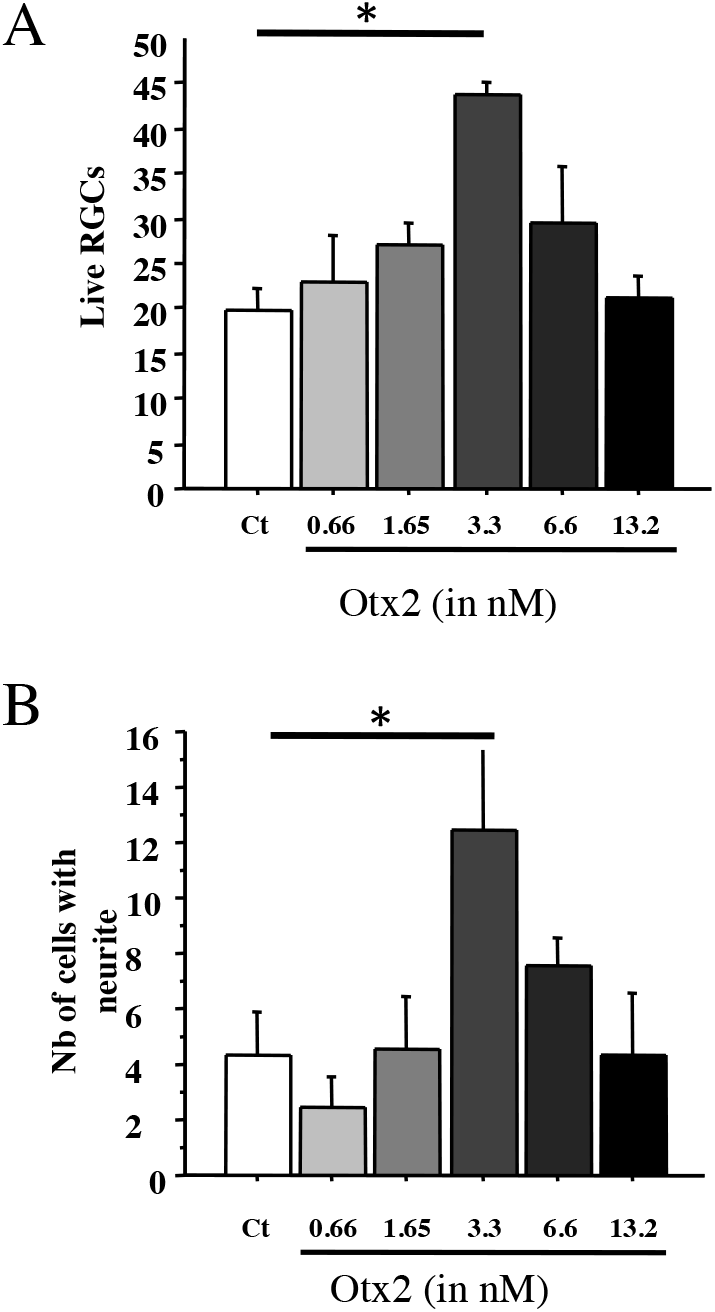
A, B: RGCs were purified from adult rat retina treated as indicated and and cultured for six days. Live cells and live cells with a neurite greater than two soma diameters were counted within a vertical coverslip diameter. Comparisons were made by one-way ANOVA followed by Fisher’s PLSD for post hoc comparisons. *p<0.05.

Retinal explants from adult rat retina grow axons that can be labelled with anti-NF200 antibodies. In cultures of vehicle treated explants, axons tended to be isolated and their cumulative length was about 9 mm in 6 days (Figure 2A,C). Explants treated with 13.2nM OTX2 grew out axons about four times longer in cumulative length (Figure 2B,C). This growth is likely to be an underestimate of total growth since the bundles were dense and formed of multiple axons. The growth of axons from explants cultured on poly-D-lysine without laminin was considerably less but OTX2 still increased cumulative axon growth in a dose-dependent manner and at 24.6nM this increase was by about 10-fold (Figure 2D).

**Figure 2.**
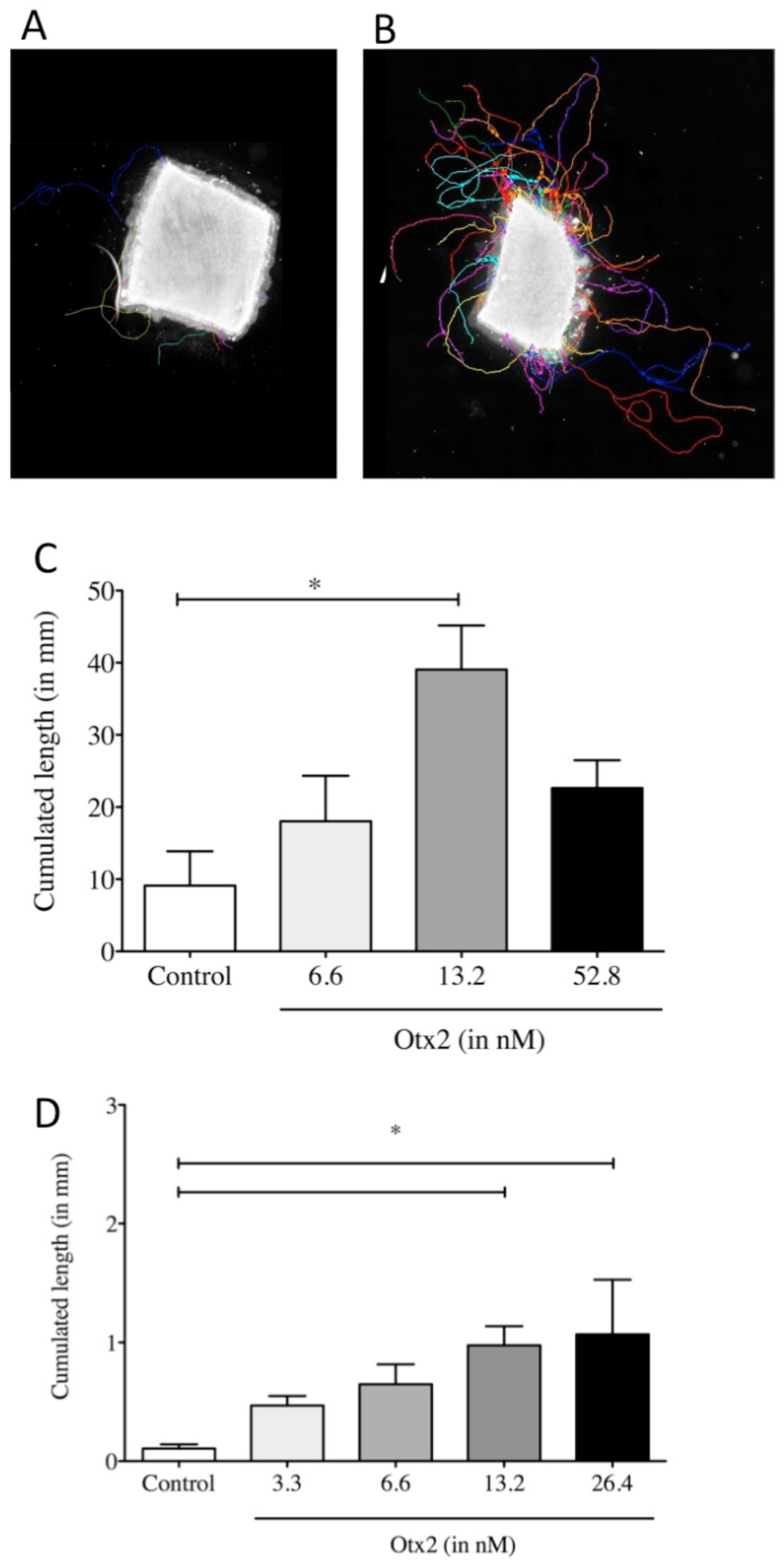
A-C: Explants from adult rat retina were treated as indicated and cultured for 6 days on laminin/poly-D-lysine-coated coverslips and then processed for NF-200 immunoreactivity. Axons and axon bundles were manually traced (colored traces) and the cumulated axon length determined. D: Cumulated axon length for explants from adult mouse cultured on poly-D-lysine without laminin and treated as indicated and cultured for six days. Comparisons were made by one-way ANOVA followed by Fisher’s PLSD for post hoc comparisons.*p<0.05

OTX2 injected soon after ONC crush increased RGC survival. ONC greatly reduced the density of Brn3a-labeled RGCs compared to an intake retina (Figure 3A-C). Retinas from mice treated with OTX2 clearly had more labeled RGCs. In terms of numbers, whereas adult C57 Bl6 mice typically have about 3300 RGCs/mm2 (Jeon et al., 1998), fourteen days after ONC, mice treated with vehicle showed a marked reduction in the number of surviving Brn3a-labeled RGCs with only about one-fifth that number (Figure 3D). In the retinas from OTX2 injected mice had about a thousand RGC/mm2, a modest (66%), but significant increase in the number of Brn3A-labeled surviving RGCs compared to vehicle (Figure 3D) and about one-third the number of RGCs in an intact retina.

**Figure 3.**
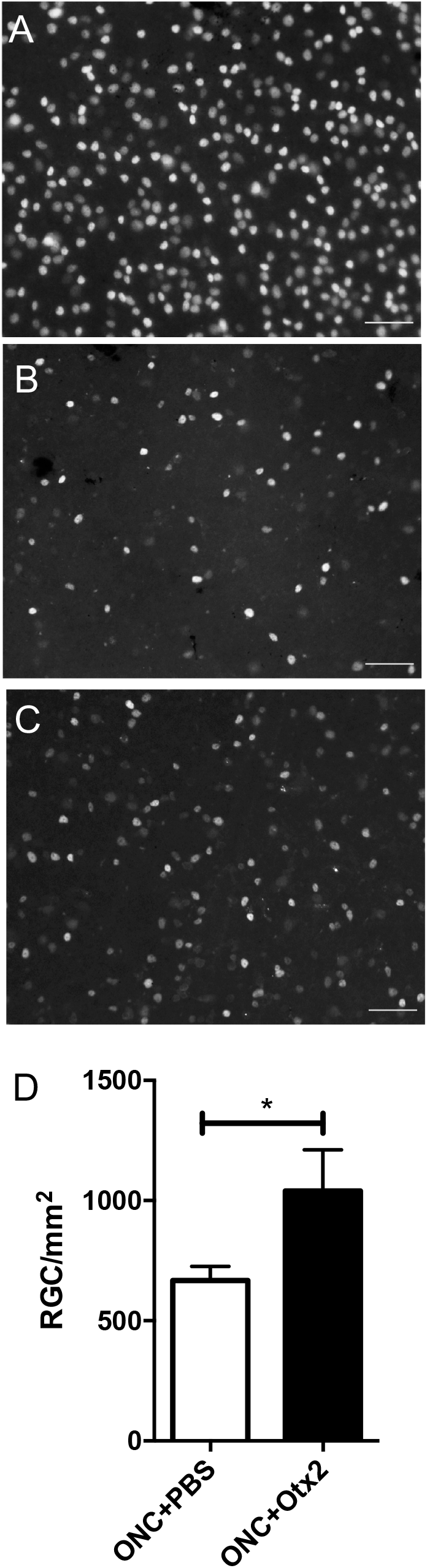
A: RGC survival two weeks after ONC. RGCs were identified by Brn3A immunofluorescence and counted in retinal flatmounts. OTX2 increased RGC survival by about 60%. P<0.05, t-test. N=14 or 15 per treatment.

OTX2 significantly increased and the number of GAP-43-positive fibers/growth cones in the distal optic nerve compared to vehicle injected mice. Longitudinal sections through the optic nerve showed showed bright GAP-43-labled regenerating axons and what might be growth cones (Fig 4A, B). At 0.5mm from the crush site the density of GAP-4-labled fibers was apparently similar (not shown). Near the optic chiasm 4.5 mm from the crush site the number of regenerating GAP-43-labled axons was strikingly different between the groups (Fig 4A, B). We counted the GAP-43-labled axons along the nerve from the crush site to the chiasm (Fig 4C)). Near the crush site nerves from mice in both conditions had similar numbers of labled fibers (95 for vehicle and 88 for OTX2). At more distal regions OTX2 treated mice had about an average of 50-65 regenerating axons at least up to the optic chiasm 4.5mm from the crush site (Figure 4C). In nerves from mice treated with vehicle, the number of GAP-43-labeled profiles diminished with distance with 20 axons at 3mm from the crush site and 9 adjacent to the chiasm, whereas mice treated with OTX2 had significantly more, 54 and 68 at the same distances (Figure 4C).

**Figure 4.**
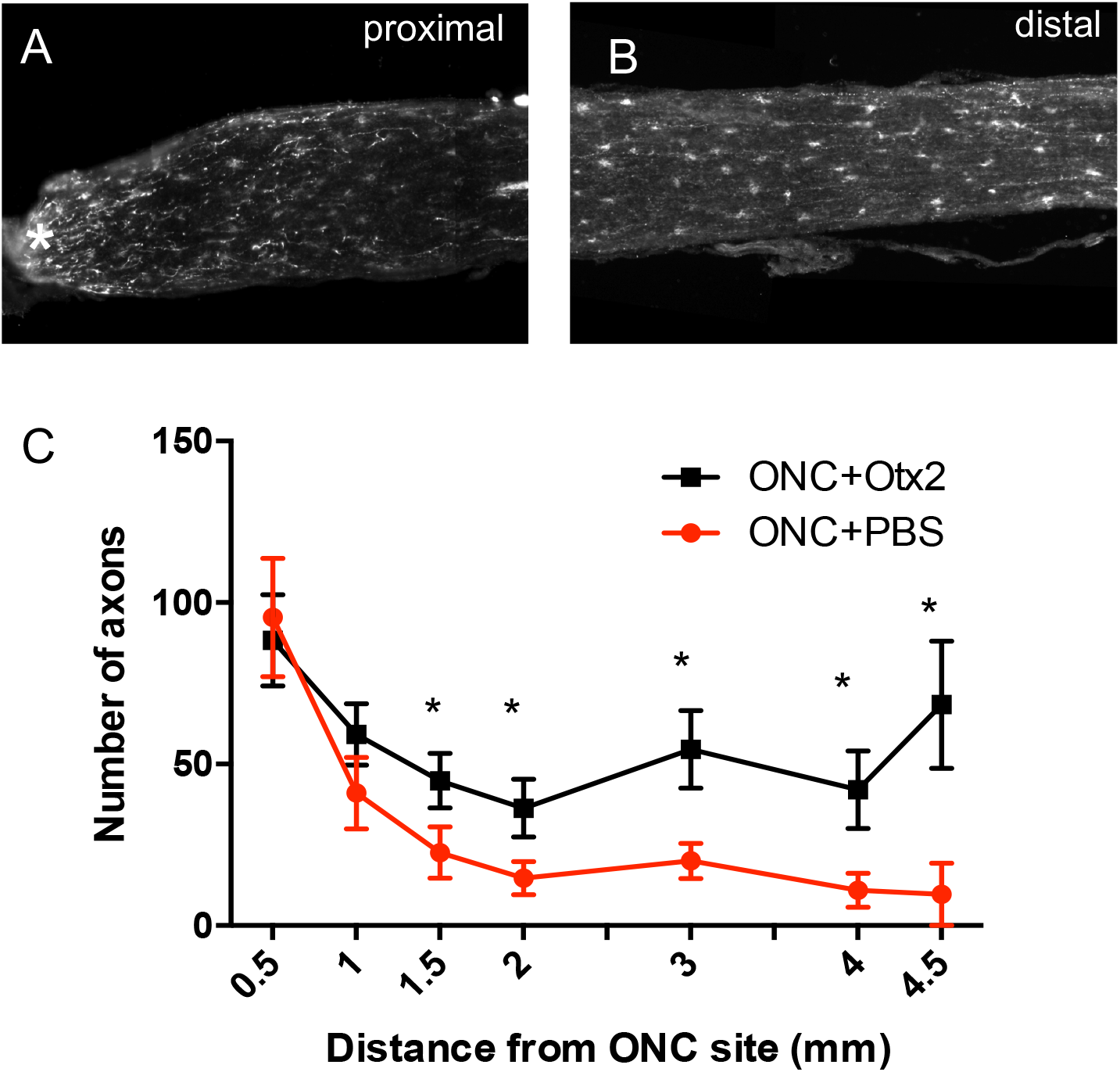
A, B: The number of GAP-43-positive regenerating axons along the optic nerve 14d after ONC proximal and distal to the crush site,. Starting at 1.5 mm from the ONC site there were significant more regenerating RGC axons in mice treated with OTX2 compared to those treated with vehicle. *p<0.05, one-tailed t-test. N=5-14 per time point per group.

We then examined whether OTX2 treatment could restore visual function. Mice were randomly assigned to two groups destined to vehicle consisting of the bacterial extract or OTX2 injection and tested for visual acuity (Figure 5). Optomotry showed that each group had a mean peak acuity of about 0.35-0.375c/deg (Figure 5). When tested three weeks post crush most mice were blind or showed highly reduced visual acuity (Figure 5; note the change in scale for the Y axis), with 10 of the mice showing no counterclockwise head movements and two responding very poorly. Thirteen weeks after crush, the left eye was sutured shut and mice tested. Four of six mice treated with Otx2 responded to the visual stimuli presented to the right eye, although visual acuity was far inferior to that measured before ONC (note the difference Y-axis in scale). None of the vehicle treated mice responded to the visual stimuli.

**Figure 5.**
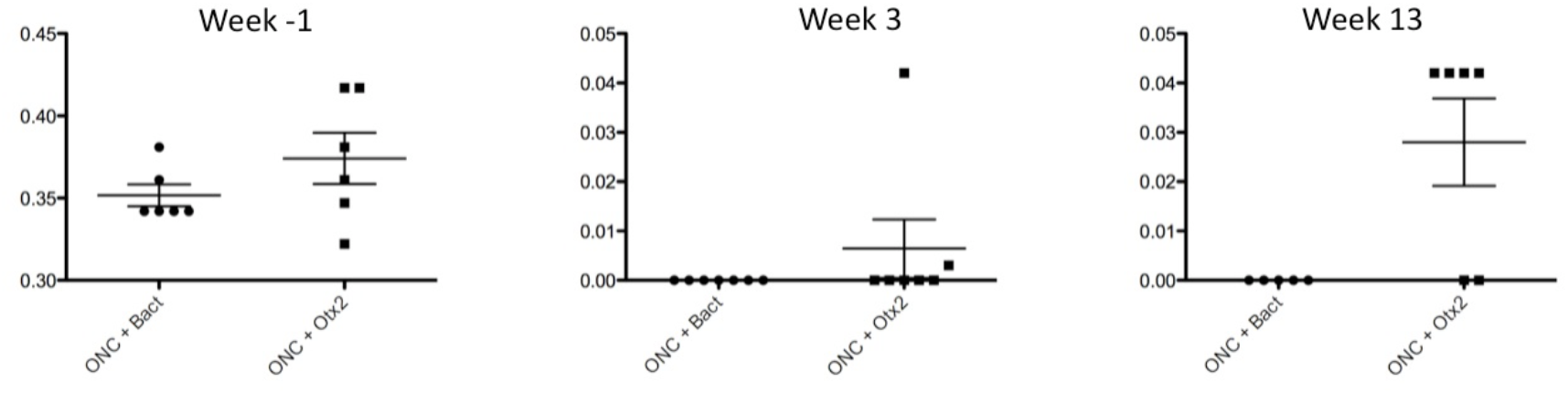
Visual acuity assessed by the optomotor test at one week before ONC, 3 weels after ONC and 13 weeks after ONC. Vertical axis is cycle/degree. Note the change in vertical axis scale between one week before and times post ONC.

**Figure 6.**
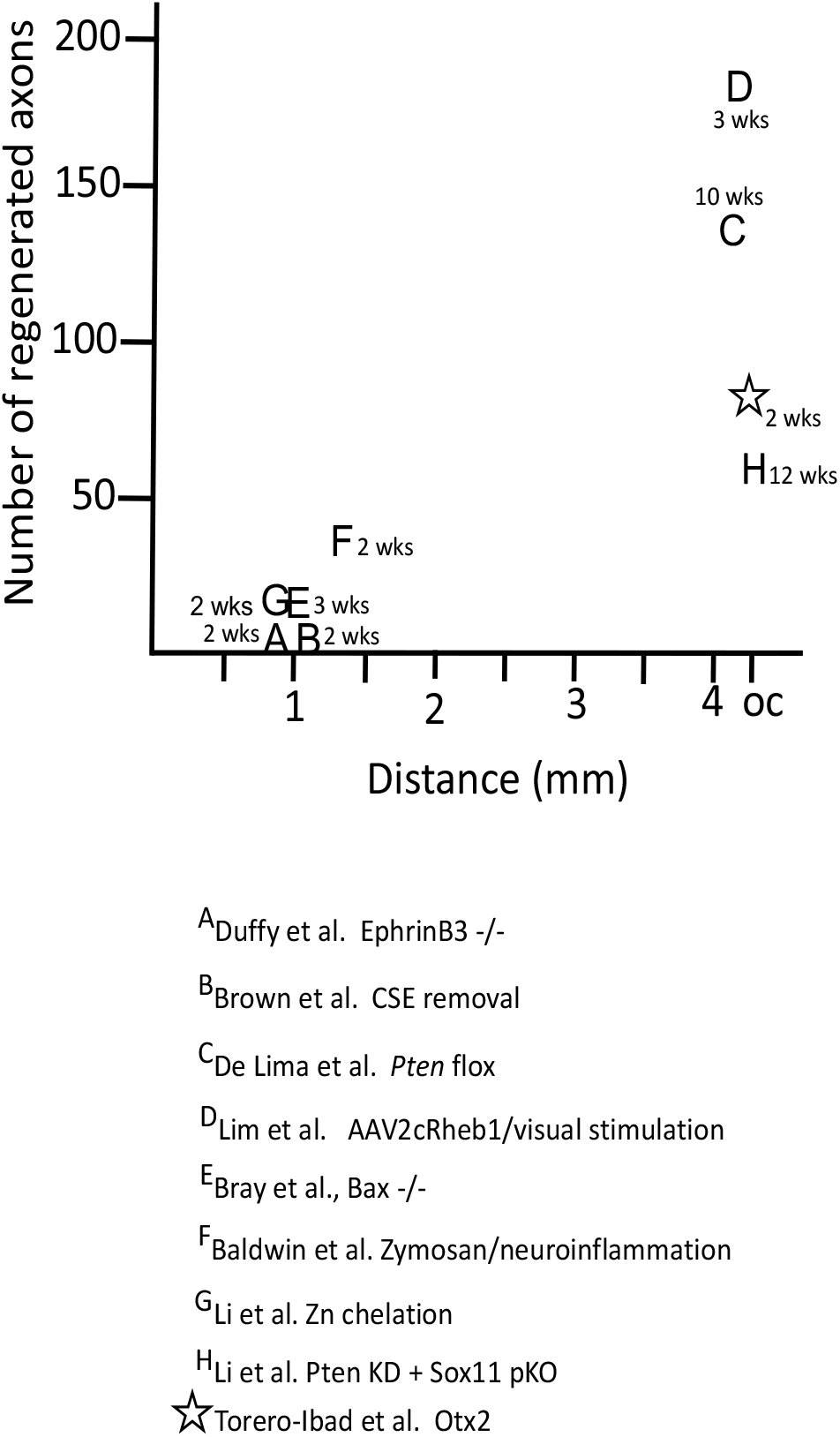
Comparison of the number of regenerating axons reported in the references and the distance. The time in weeks is the time for the axons to reach longest endpoint.

## Discussion

Without manipulation, mammalian adult CNS axons including those of RGCs fail to successfully regenerate after axotomy. An important objective in neurobiology has long been to find conditions, factors or treatments that stimulate adult axon regeneration and restore CNS function. Here we provide evidence that the homeoprotein transcription factor OTX2 stimulates RGC axon regeneration *in vitro* and *in vivo* in two rodent species.

Dissociation of the retina is a harsh treatment that shears off RGC dendrites and axons and leads to the death of almost all cells. Here, we independently corroborate previous findings that exogenous OTX2 promotes RGC survival and now demonstrate that it stimulates axon regeneration. The fact that the fibers are NF 200 positive indicates that they are axons. Regarding RGC survival, we observed bell-shaped dose-response curve similar to that already reported (Torero Ibad et al., 2011). The dose of exogenous OTX2 that stimulated RGC axon regeneration in mixed cultures was higher than that promoting RGC survival, suggesting a possible inhibitory effect of the other cell types present and demonstrating that cell survival does not necessarily imply a neurite outgrowth. An inhibition of neurite outgrowth by the non-RGCs cells is supported by the fact that in a situation where adult RGCs have been purified, thus respond directly to OTX2 for their survival (Figure 1C and (Torero Ibad et al., 2011)), the dose-response curves for cell-survival and neurite outgrowth are almost identical.

Retinal explants avoid separating RGC cell bodies from their proximal axons and dendrites and retinal cells remain in their tissue context at least initially. In control conditions, few axons grew out from the edges of the explants from adult retinas during the 6 days of the culture. However, OTX2 at doses higher than those that promote RGC survival significantly stimulated RGC axon regeneration. Adult mouse explant cultures on laminin/poly-D-lysine grew out axons with a cumulative length of at least 40 mm in six days (Fig 2A-C). Even on a substrate that poorly supports axon regrowth (i.e., poly-D-lysine alone) OXT2 still stimulated significant growth.

Optic nerve crush leads to massive RGC death and axon loss. Administration of OTX2 5 to 15 minutes after ONC provides a modest but significant increase (about 60%) in RGC survival (Figure 3D). Previously it was shown we that OTX2 protected RGCs against NMDA excitotoxicity *in vivo* and the present results add a second model for *in vivo* RGC neuroprotection by OTX2. More importantly, OTX2 greatly increased the number of GAP-43-labeled regenerating axons in the distal nerve (Figure 4C). In mice treated with vehicle after crush up to the optic chiasm only 9 GAP-43-labled profiles were observed in the distal nerve 14 days after ONC. In contrast, on average 68 regenerating axons reach the optic chiasma in mice treated with OTX2 (Figure 4D).

OTX2 appeared to partially restore visual acuity to a very low level after ONC (Figure 4). Before ONC mice had a visual acuity threshold of about 0.375 c/deg. Three weeks after nerve crush, they did not respond to the visual stimulus. After 13 weeks four out of six mice treated with OTX2 responded to the visual stimuli but at very low visual acuity of about 0.03 c/deg, while none of the vehicle treated mice responded to the stimuli. Since the left eye was sutured closed we are sure that the responses to the visual stimuli, albeit very weak, were driven by the OTX2-treated eye. A limitation of this experiment is that we were unable to visualize regenerated axons. Cholera toxin βsubunit anterograde tracing did not work and 13 weeks is too late after the elongation period of axon growth for the axons to be labeled with GAP-43 (Moya et al., 1988). Thus, in this experiment we were unable to quantify regeneration or to determine how far past the chiasm any regenerated fibers reached.

While Otx2 stimulates RGC axon regeneration in vitro and in vivo, is the amount of regeneration meaningful? C57 Blk6 mice have about 50-52000 RGC axons in the optic nerve (Jeon et al., 1998; Templeton et al., 2014) and these 50-52000 RGCs and axons underpin all visual behaviors, some of them complex. A first important issue is how does OTX2 stimulate regeneration compared with other approaches. Figure 5 compares the number of regenerating axons at their maximal distance after ONC crush in different contexts. Many strategies resulted in well less than 50 axons regenerating to 1-1.5mm from the ONC site. (Duffy et al., 2012) observed 10 or so axons reaching 1mm using EphrinB3 knockout mice. (Brown et al., 2012) reported similar results in mice in which chondroitin sulfate E was removed. There are only three previous reports of RGC regeneration reaching the chiasm and only one in the time window that we observed here. De Lima (2012) (de Lima et al., 2012) reported that about 150 axons reach the chiasm 10-12 weeks after ONC after floxing *Pten*. (Li et al., 2017) used zinc chelation to stimulate regeneration of about 60 axons to the chiasm after 12 weeks. And (Lim et al., 2016) reported about 180 regenerating axons at the chiasm three weeks after crush after expressing a positive regulator of mTOR signaling combined with high contrast visual stimulation. These approaches are not easily translatable for use in humans. Here we observed about 70 regenerating axons at the chiasm two weeks after ONC with OTX2 treatment.

Other recent studies manipulating inflammation (Baldwin et al., 2015); using transgenic mice resistant to apoptosis (Bray et al., 2019), altering retinal zinc (Li et al., 2017) or combining Pten knockdown with Sox11 knockdown (Li et al., 2018) or overexpression of Lin28 (Wang et al., 2018; Zhang et al., 2019) report regeneration success rates in terms of number of regenerating axons at the chiasm 2 weeks post crush inferior to our results. An important difference in our approach is that we provided additional transcription factor that is already present in the retina without any genetic manipulations.

A second issue is whether the average of 50, best case of 205 regenerated RGC axons could provide the substrate for the apparent slight recovery of vision we observe with Otx2. About 120 axons reaching the chiasm at 10 weeks post crush or about 180 axons at the chiasm at 3 weeks have been reported to be sufficient for visual behaviors such as responding to the visual cliff, optokinetic response, papillary light reflex, photoperiodicity and looming response (de Lima et al., 2012; Lim et al., 2016). Thus, although 50-205 axons out of some 50000 appears to be a minuscule fraction of the intact visual pathway, Otx2 treatment may provide results similar to what was reported in other studies reporting functional recovery.

How might Otx2 increase adult RGC regrowth after injury? One possibility is that the increased axon regeneration may reflect the increase in RGC survival (see Figure 3A). In cat retina, adult alpha cells are more resistant to axotomy and appear to be more capable of at least short distance regeneration (Watanabe et al., 1995). The present results could be explained if among the RGCs spared death by Otx2 there are cells corresponding to those resistant to axotomy and/or Otx2 had a preferential effect on regeneration competent cells. In terms of molecular mechanism, Lin28 is of interest because it is expressed in amacrine cells and during development its expression is regulated by OTX2 (Parisi et al., 2017; Zhang et al., 2019). OTX2 is not expressed in amacrine cells of the adult GCL, but all cells including displaced amacrine cells can take up exogenous OTX2 (Torero Ibad et al., 2011). Thus, it is conceivable that OTX2 injected into the eye and taken up by amacrine cells could influence expression of Lin28 and stimulate regeneration. Future studies are needed to explore this possibility.

In addition to promoting significant RGC axon regeneration, OTX2 also promotes the survival of adult RGCs in vitro and in vivo ((Torero Ibad et al., 2011) and present results). This might be useful in the context of glaucoma, diabetic neuropathies or neuromyelitis optica. These diseases have been considered axonopathies in which RGC axons start to degenerate due to constriction at the optic head or by autoimmune attack before frank RGC loss (Calkins, 2012; Casson, 2012; Kern and Barber, 2008). In patients diagnosed with such diseases, OTX2 could promote survival of the RGCs, long enough to allow regeneration by OTX2 alone or in combination with another regeneration stimulating approach. One advantage of OTX2 as a possible therapeutic agent is the fact that it is already present in the retina and injecting it into the eye is simply adding more to what is already present. This obviates the need for gene therapy, cell therapy, or administering compounds that are not endogenous to the human body. Furthermore, the ability of exogenous OTX2 to enter RGCs (Torero Ibad et al., 2011) eliminates the need for vectors or targeting strategies.

## Acknowledgements

We thank J. Dégardin and M. Simmonuti of the Institut de la Vision, Paris for visual acuity testing and Prof. Larry Benowitz and Dr. Yiqing Li of Boston Children’s Hospital for assistance in preliminary ONC experiments. This work was supported by Fovea-Pharmaceuticals, and Global Research Laboratory Program Grant 2009-00424 from the Korean Ministry of Education, Science, and Technology, ANR NeuroprOTX, ERC

